# Development of a core genome Multilocus Sequence Typing (cgMLST) and Life Identification Number (LIN) code schema for the *Burkholderia cepacia* complex

**DOI:** 10.64898/2026.11.10.765358

**Authors:** Kasia M Parfitt, Keith A Jolley, Charlotte Peeters, Made Ananda Krisna, Eshwar Mahenthiralingam, Martin CJ Maiden

## Abstract

**Background:** The *Burkholderia cepacia* complex (Bcc) currently comprises over 20 closely related species; however, understanding Bcc population biology is complicated due to the continual advances in taxonomy and diversity driven by extensive horizontal gene transfer (HGT). Whilst there are conventional typing schemes for the Bcc, including Multilocus Sequence Typing (MLST), there remains a gap for fine-scale resolution typing of the species complex. Here, we present a 1,925 loci core genome MLST (cgMLST) and 14-digit Life Identification Number (LIN) barcoding scheme for the Bcc, that is embedded in the PubMLST database (https://PubMLST.org/organisms/burkholderia-cepacia-complex).

**Methods:** A dataset of 142 complete Bcc genomes spanning 19 species were downloaded from NCBI to develop the draft cgMLST scheme (Dataset 1). Two pangenome tools (PIRATE and PEPPAN) were then combined to give a consensus core gene list from Dataset 1. The ‘core genes’ from these outputs were then screened against the PubMLST loci, using a sequence tag scan, to identify their presence and curate new alleles. The core genes were then validated using a larger Bcc dataset (*n* = 934 genomes; 22 species) on PubMLST, which had been filtered for high-quality (≤200 contigs and 100% ribosomal MLST confidence). This refined the core gene list, ensuring a robust and reliable scheme using a threshold of 99% to classify the gene as ‘core’. The population structure of the Bcc was subsequently analysed using the validation dataset to identify genetic clusters using ReporTree. Combined with statistical analysis, 14 allelic mismatch thresholds were identified for the LIN codes.

**Results:** The Bcc cgMLST scheme consists of 1,925 loci, which have been designated an arbitrary ‘BCC’ identifier. The cgMLST scheme and 14-threshold LIN code has been implemented in PubMLST and are publicly available for use. The Bcc genomes in the PubMLST database have been assigned cgSTs and LIN codes. Use cases of the LIN codes were validated using three previously published datasets, which spanned both outbreak investigation and phylogenomic lineage structures.

**Conclusions:** This stable, hierarchical LIN code nomenclature enhanced the ability to cluster isolates at multiple levels of resolution with examples overlayed on published datasets provided. The LIN code system is also backwards compatible with previous MLST and clonal complex (CC) nomenclature. Overall, the updated typing scheme will facilitate improve global public health through surveillance of the Bcc epidemiology including source tracing and outbreak investigation.

## Introduction

The *Burkholderia cepacia* complex (Bcc) is a closely related group of environmental bacteria frequently found in clinical settings (Baldwin *et al*., 2005), which comprises 24 named species (Morales-Ruíz *et al*., 2022) at the time of writing (July 2026). Two additional species, “*Burkholderia paludis”* (Ong et al., 2016) and “*Burkholderia reimsis”* (Esmaeel et al., 2018), have been proposed but are not validly published nomenclatures (Morales-Ruíz *et al*., 2022). The complex comprises both variants that are opportunistic pathogens which cause infection (Mahenthiralingam *et al*., 2005) and those of benefit to the environment and agriculture (Parke and Gurian-Sherman, 2001). Bcc pathogens are especially problematic to immunocompromised individuals, such as those with cystic fibrosis (CF) or chronic granulomatous disease (CGD) (Mahenthiralingam *et al*., 2005). Individuals with CF may experience ‘cepacia syndrome’ from severe lung decline (Ledson *et al*., 1998; Parfitt *et al*., 2023). Less is known about the interaction between the Bcc and individuals with CGD (Zelazny *et al*., 2009); although, association has been made with pneumonia (Greenberg *et al*., 2009; Maignan *et al*., 2013). Bcc bacteria are a common cause of contamination in sterile health care (Tavares 2020) and non-sterile industrial products (Cunningham-Oakes *et al*., 2021), both of which may lead to outbreaks of infection. The Bcc is of clinical importance due to their intrinsic resistance to antibiotics (Aaron *et al*., 2000), which contributes to difficulties in treatment.

Bcc infections are not linked to commensal carriage, and it is thought that Bcc infection arises in hospital settings or the environment (Tavares *et al*., 2020). Previous evidence has detailed patient-to-patient spread among CF individuals within the Bcc in hospital settings (LiPuma *et al*., 1990; Govan *et al*., 1993). Due to stringent segregation methods for infection control of individuals with CF in the 1990s (Saiman *et al*., 2000), patient-to-patient spread diminished (France *et al*., 2008). However, novel sequence types (STs) continue to emerge and cause CF infection which are thought to be from environmental reservoirs. For example, *B. multivorans* has been shown to have limited within-patient evolution but high genomic diversity between patients (Lood *et al*., 2021). Chronic Bcc infection has also occurred among siblings with CF, outside of the hospital setting (Mahenthiralingam *et al*., 2005). During chronic infection, different Bcc species out-compete each other in the CF lung environment (Mahenthiralingam *et al*., 2001). In addition to the importance of the Bcc in immunocompromised infection, this group is also implicated in numerous industrial– (Cunningham-Oakes *et al*., 2021) and hospital-related outbreaks (Spilker *et al*., 2022) including contamination of medicinal products (Tavares *et al*., 2020), hygiene products (Lo Cascio *et al*., 2006; Molina-Cabrillana *et al*., 2006; Bender *et al*., 2022), and medical equipment (Nannini *et al*., 2015; Doran *et al*., 2025). The ability of the Bcc to colonise an array of niches, both natural and man-made, is consequence of genotypic and phenotypic plasticity and ability to survive in nutrient-scarce conditions (Tavares *et al*., 2020). Therefore, it is important to understand and control the presence of the Bcc in such environments as they pose a significant risk to immunocompromised individuals.

Classification of the *Burkholderia* genus has developed over time, including the identification of new species, and the reclassification of historical taxonomic groupings. At the time of writing, there were 24 named species in the Bcc (Jin *et al*., 2020). Accurate identification and classification of the Bcc is crucial for understanding its epidemiology, risks for CF lung infection, and outbreak investigations. Classical microbiological isolation and identification methods are widely available and low-cost, but do not provide sufficient resolution among Bcc species (Coenye *et al*., 2001; Vanlaere *et al*., 2009; Silva-Santana *et al*., 2026). Identification of the Bcc remains challenging due to the high similarity among Bcc species phenotypically and genetically (Silva-Santana *et al*., 2026). The use of molecular nucleotide sequence-based methods for Bcc identification and differentiation have proven effective, including the use of the *recA* gene (Mahenthiralingam *et al*., 2000). In 2005, sequence-based analysis was expanded to include a Multi Locus Sequence Typing (MLST) scheme using 7 housekeeping genes for more accurate Bcc differentiation (Baldwin *et al*., 2005). The original MLST scheme was subsequently expanded by the development of PCR primers to differentiate between Bcc species and isolates, amplifying target loci from multiple species within *Burkholderia* sensu lato (Spilker *et al*., 2009). Whilst MLST provides a good platform for Bcc species identification, with discriminatory power for basic pathogen epidemiology (Maiden *et al*., 2013) due to the complex’s low frequency of point mutations (Waine *et al*., 2007), there remain gaps in how it can be applied. For instance, greater resolution is required for source tracking in the Bcc. This was illustrated in a publication looking at *B. multivorans* ST-742 in Belgium, where multiple patients were shown to harbour this ST but without evidence for cross-infection (Lood *et al*., 2021). Source tracking is needed to identify transmission routes and pinpoint the environmental sources to connect human isolates to specific reservoirs (i.e. medical equipment, contaminated products, and pharmaceuticals). Additionally, the identification of persistent or recurrent sources can be identified by comparing isolates from different temporal backgrounds to the same reservoir.

With the advent of whole genome sequencing (WGS), greater numbers of Bcc genomes are available (>1,900 genomes within PubMLST as of October 2026). This enhances the capability to define evolutionary relationships, perform comparative genomics, and differentiate lineages within Bcc species (Jin *et al*., 2020; Parfitt *et al*., 2023) and across *Burkholderia* sensu lato (Mullins and Mahenthiralingam, 2021). By utilising WGS data, we can expand 7-locus MLST into a core genome MLST (cgMLST) scheme, which uses thousands of genes, to enhance the precision of detecting genomic variants (Maiden *et al*., 2013; Schürch *et al*., 2018). As homologous recombination can occur amongst isolates and species of the Bcc, mediated by insertion sequences (IS) and genomic islands (Waine *et al*., 2007; Baldwin *et al*., 2008; Silva-Santana *et al*., 2026), cgMLST is advantageous as it dilutes recombinogenic effects (Preska Steinberg *et al*., 2022; Krisna *et al*., 2024). Coupled with cgMLST as a foundation, Life Identification Number (LIN) codes (Palma *et al*., 2026) can be developed to provide a stable, multi-level nomenclature approach for the Bcc complex. For the application of LIN codes, thresholds are chosen using a pragmatic approach, identifying discontinuities in the bacterial population. LIN codes provide a unique genomic barcode to each genome, which is automatically assigned using a database such as PubMLST (https://pubmlst.org) (Jolley *et al*., 2018). The barcodes are stable as new genomes are introduced into the database and ultimately provide a rapid visualisation of genetically similar isolates to the user to facilitate public health surveillance of emerging lineages.

We present a cgMLST and LIN code genomic framework for the Bcc for the use in epidemiological studies and surveillance, including source tracking. To do this we validated a cgMLST scheme and LIN code nomenclature for studying the population biology of the 24 Bcc complex species defined at the time of writing. This Bcc-combined scheme provides a common genomic framework for the comparison of biologically similar isolates with better handling in the instance of uncertain/changing taxonomy. By creating the scheme in a broad manner, we will also enable future research around the genomic relationships and epidemiological clusters of different species within the Bcc due to their occupation of the same niches. Both the cgMLST and LIN code schemes are publicly available and can be used globally to characterise Bcc genomes in a portable and standardised manner.

## Methods

A workflow was developed for the Bcc cgMLST and LIN code development and validation (Figure 1).

**Figure 1.**
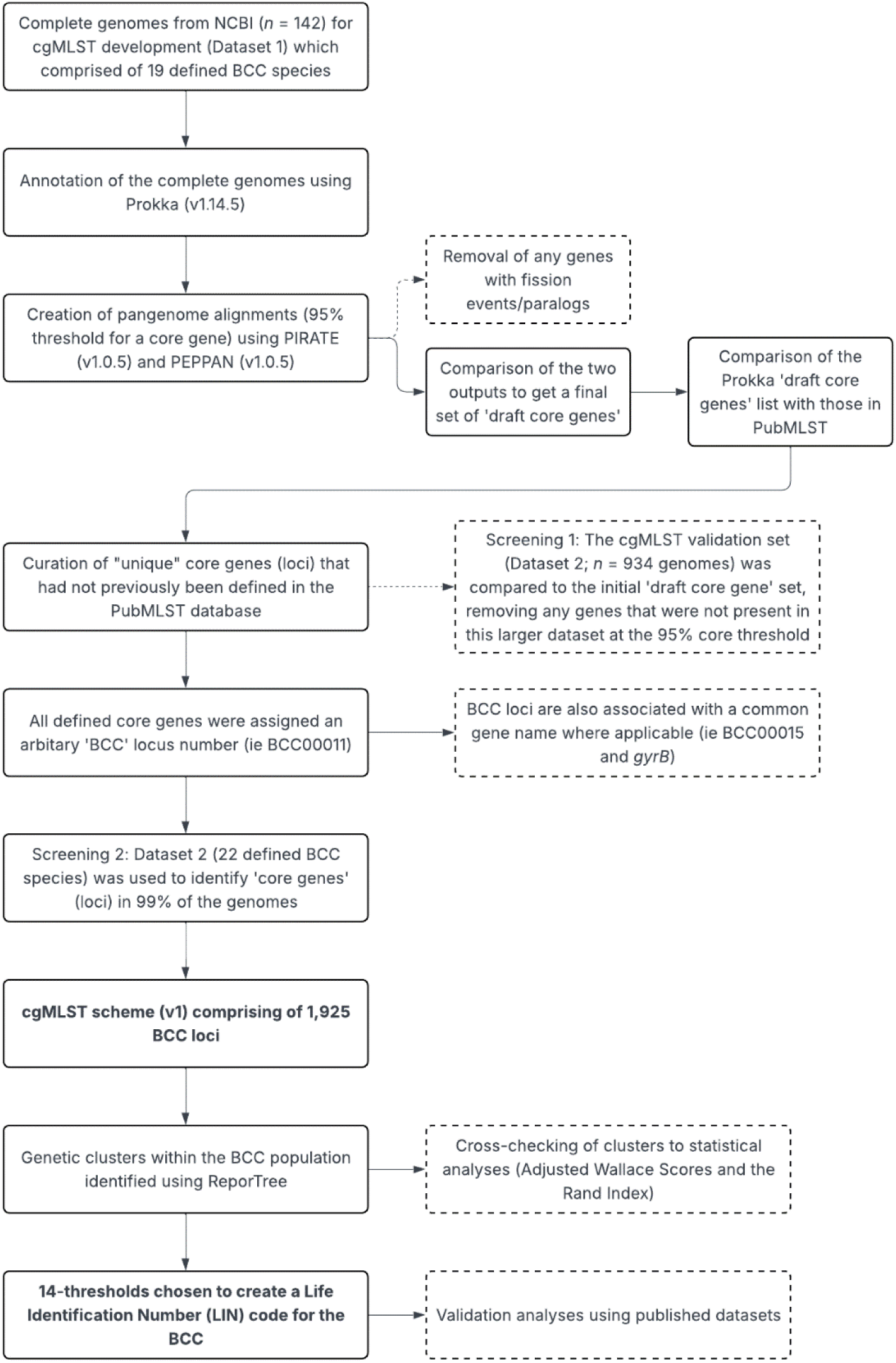
| The workflow describing BCC cgMLST and LIN Code schema development and validation.

### Dataset compilation

The datasets used in this study were compiled from either (i) National Center for Biotechnology Information (NCBI) (https://www.ncbi.nlm.nih.gov) for the development dataset; or (ii) the Bcc database on PubMLST (https://pubmlst.org/organisms/burkholderia-cepacia-complex) for the validation dataset (Jolley *et al*., 2018). On 1^st^ August 2023, all complete Bcc genomes on NCBI (*n* = 162) were downloaded and uploaded to PubMLST; a database that contains (i) isolate metadata, (ii) genome sequences, and (iii) MLST profiles and clonal complexes. The 162 genomes were filtered for a ‘good’ ribosomal MLST (rMLST) score (defined as *n =* 53; 100% loci designated) and cross-checked for matching species identification against the NCBI deposits. After filtering, the developmental dataset (known as Dataset 1) comprised of 142 Bcc genomes and represented 19 Bcc species.

A larger dataset was chosen from PubMLST (accessed on 1^st^ August 2024) to represent the Bcc population for the cgMLST validation analyses (known as Dataset 2). This dataset was also used to develop the LIN code schema. All isolates in the database at this time (*n* = 1,119 genomes) were assessed for species identification using the rMLST designation outputs (only using those with 100% confidence) and filtered for ‘high quality’ draft genomes that had ≤200 contigs. After filtering, there were 934 genomes that were used for validation purposes (Dataset 2). This dataset represented 22 defined Bcc species (plus the group ‘other Bcc’, as defined in PubMLST at the time). Isolates were from 31 countries across all 6 continents (excluding Antarctica). Dataset 2 also encompassed 6 different sources (CF patient, non-CF patient, hospital environment, environmental, industrial, CGD patient) to provide a global distribution.

### Updating of clonal complexes (ccs) in the PubMLST database

All STs in the PubMLST database at the time of writing (*n* = 2,517) (26^th^ January 2026) were downloaded and assessed for clonal complexes (ccs). It was identified that some previously assigned ccs did not have (i) a central ST, or (ii) enough isolates in the database for cc assignment. The STs were subject to analysis with the Phyloviz Desktop software (v.2.0a) (Nascimento *et al*., 2016). A full MST goeBURST was performed to identify clustering groups (Supplementary Figure 1). These were then cross-checked against the PubMLST (Jolley *et al*., 2018) typing database (https://pubmlst.org/bigsdb?db=pubmlst_bcc_seqdef) using the BURST plugin. Groups that had a central ST and ≥10 isolates in the database were assigned a respective cc (Supplementary Table 1).

### Core gene identification, validation and curation

Dataset 1 (*n* = 142 genomes) was annotated for protein-coding genes using Prokka (v1.14.5) (Seemann, 2014). Two pan genome pipeline analysis tools, PIRATE (v1.0.5) (Bayliss *et al*., 2019) and PEPPAN (v1.0.5) (Zhou *et al*., 2020) were used to assess the core and accessory genome of the 142 Bcc genomes, using the GFF output files from Prokka (v1.14.5) (Seemann, 2014) as an input. For the generation of an initial set of ‘core genes’, a threshold of gene presence in 95% of isolates or greater was used in both pipelines. PIRATE (v1.0.5) (Bayliss *et al*., 2019) was ran with the –a and –r flags to align all genes and produce core/pangenome alignments, and plot summaries using R, respectively. PEPPAN (v1.0.5) (Zhou *et al*., 2020) was ran using –-match_identity 0.7 for a minimum identity of 70% in the BLAST (Altschul *et al*., 1990; Camacho *et al*., 2009) searches. PEPPAN_parser (Zhou *et al*., 2020) was subsequently run using the –t, –c flags to generate a gene presence/absence tree and refraction curve, respectively. Any genes that presented putative paralogs/fission events were excluded from the datasets prior to comparing the two outputs. Removal of these genes were important to ensure automated annotation of core genes from Bcc genomes deposited into PubMLST (Jolley *et al*., 2018). The ‘draft core genome’ was created based on the remaining genes that were consistent between both the PIRATE (Bayliss *et al*., 2019) and PEPPAN (Zhou *et al*., 2020) outputs.

The complete genome of *B. ambifaria* Q53 (Nievas *et al*., 2023) was used as a recently sequenced reference to initially define Bcc loci in the PubMLST database. The ‘draft core genes’ were cross-referenced to loci in the PubMLST Bcc database (https://pubmlst.org/organisms/burkholderia-cepacia-complex) (Jolley *et al*., 2018) for curation, curating new alleles where applicable. Dataset 2 (*n* = 934 genomes) was used to validate the ‘draft core genes’ using the BIGSdb Genome Comparator plugin on PubMLST (Jolley *et al*., 2018) at a 95% core gene threshold. Any genes that were absent in Dataset 2 compared to Dataset 1 were removed from the analysis. Unique alleles, that had not previously been identified in the database, were manually curated using the ‘sequence tag scan’ option with default parameters (minimum percentage identity of 70%, minimum sequence alignment of 50%, and a BLASTn word size of 20). After curation, a ‘final core gene’ list was obtained using a 99% core threshold, to ensure a robust cgMLST schema. All defined core genes were assigned an identifier that comprised of a ‘Bcc’ prefix, followed by an arbitrary locus number. Where applicable, these Bcc locus numbers can be associated with a common gene name (i.e. Bcc00015 with *gyrB*).

The threshold of allowed missing loci was defined as 38 (2% loci) within the Bcc cgMLST scheme. Those that met this threshold were assigned a core genome ST (cgST) using the automated profile definer on BIGSdb (https://github.com/kjolley/BIGSdb/blob/develop/scripts/automation/define_profiles.pl). Genomes are provided with categories based on their cgMLST assignment scores as follows: (i) ‘good’ is genomes with fewer than 38 missing loci, (ii) ‘intermediate’ are those with between 38 and 100 missing loci, and (iii) ‘bad’ as those with over 100 missing loci.

### Assessment of allelic variability and gene COG analysis

The Bcc cgMLST loci were assessed for the total number of allelic variants and their respective gene lengths using a previously published Python script (https://github.com/artmisk13/cgmlst_hinf/blob/code/core_gene_allelic_variability.py) (Krisna *et al*., 2024). Annotation of the core genes was then assigned through eggNOG-mapper (v2.1.13) utilising the Diamond database in blastx mode with default parameters (Cantalapiedra *et al*., 2021). A putative Cluster of Orthologous Genes (COG) category (Cantalapiedra *et al*., 2021) was provided for each gene in the output. Genes that were not defined a category (output of ‘-’) were assigned to the ‘unknown function’ category ‘S’. The COG categories were further defined into the respective Kyoto Encyclopedia of Genes and Genomes (KEGG) groups (Kanehisa and Goto, 2000; Kanehisa *et al*., 2016).

### Analysis of the Bcc population to define allelic mismatch thresholds

Genetic clusters within the BCC population of Dataset 2 (*n* = 934 genomes) were identified using ReporTree (Mixão *et al*., 2023) on the PubMLST database (Jolley *et al*., 2018). All parameters were set to default, with additional metadata added for species and clonal complex. To identify the optimal genetic clusters within the BCC population, the stable regions and metrics outputs from ReporTree (Mixão *et al*., 2023) were interrogated. Stable regions that had a corresponding RandIndex score and adjustedWallace coefficient of =>0.9 were used. Thresholds were then assessed using the Grapetree (Zhou *et al*., 2018) plugin on PubMLST (Jolley *et al*., 2018) to analyse the hierarchical clustering at each level.

A 14-threshold LIN code system was established for the Bcc population. The LIN code reads from left to right and increases in the discriminatory values between isolates as the allelic mismatches decrease. The first 8 thresholds were identified using the regions of cluster stability (Figure 3). An additional 6 thresholds were pragmatically chosen to facilitate epidemiological analysis such as outbreak investigation and transmission events.

### Phylogenetic analysis

Dataset 2 (*n* = 934 genomes) was used to reconstruct phylogenetic trees for validation. The Genome Comparator plugin on PubMLST (Jolley *et al*., 2018) was used to generate a FASTA file alignment of all cgMLST loci using MUSCLE (Edgar, 2004). SNP-sites (v2.5.1) (Page *et al*., 2016) was used to identify the constant sites (-C) from the alignment for use in phylogenetic tree building. A maximum likelihood phylogenetic tree was built using IQ-TREE (v2.2.2.6) (Minh *et al*., 2020), using the GTR+F+G model with 1,000 bootstraps. The –fconst flag was used to correct for bias through providing the constant regions output from SNP-sites (Page *et al*., 2016). ClonalFrameML (v1.12) (Didelot and Wilson, 2015) was used to adjust the phylogenetic output, accounting for recombination events. The phylogenetic tree was then annotated with the respective metadata: (i) clonal complex and (ii) core gene cluster (cgc)1750. Figtree (v1.4.4) (Rambaut, 2018) was used to create subtrees of the *B. cenocepacia* ST-31 complex for in-depth analysis of the cgcs at higher resolution thresholds.

Three previously published datasets (Table 1) were used to validate the LIN codes. Maximum likelihood phylogenetic trees using the GTRGAMMA model, supported by 100 bootstraps were generated using RAxML (v8.2.12) (Stamatakis, 2014). All phylogenies were adjusted for recombination using ClonalFrameML (v1.12) (Didelot and Wilson, 2015) as above. All maximum likelihood phylogenetic trees were linked and annotated with associated metadata using the ggtree (v4.0.1) (Yu *et al*., 2017; Yu, 2020) in RStudio (v4.5.2) (RStudio Team, 2020). Final figures were assessed, assembled, and annotated using Inkscape (v1.4.4) (Inkscape Team, 2023).

**Table 1.** | Previously published datasets used for validation of the LIN codes.

| Dataset | Species | Reference |
|---|---|---|
| Healthcare-associated outbreak in Germany | *B. arboris* and *B. cepacia* | Bender *et al.,* 2022 |
| Endophthalmitis outbreak traced to clinic ventilation systems | *B. contaminans* | Spilker *et al.,* 2021 |
| Phylogenomic analysis and model isolates | *B. multivorans* | Parfitt *et al.,* 2023 |

A minimum-spanning tree (MST) was created using the core genome allelic profile and annotated with the 1^st^ LIN threshold. This utilised the GrapeTree plugin on the PubMLST database (Jolley *et al*., 2018).

## Results

### Isolate provenance information and quality statistics

The isolates used in the cgMLST development (Dataset 1, *n* = 142 genomes) were made up of 19 Bcc species, predominantly *B. multivorans* (*n* = 51, 35.9% genomes) and *B. cenocepacia* (*n* = 21, 14.8% genomes). The isolates spanned 19 countries, with most isolates (*n* = 34, 23.94%) from Germany linking to a health-care outbreak (Table 1) (Bender *et al*., 2022). Overall, the predominant isolate source was from Europe, comprising 48/142 (33.80%) of isolates. South America had the lowest isolate prevalence in the dataset with 3 genomes (2.11%) (Supplementary Table 2). The cgMLST validation isolates (Dataset 2, *n* = 934 genomes) had a greater number of species inclusion (*n* = 23) with the addition of *B. arboris*, *B. diffusa*, “*B. paludis”* and “other Bcc”. The two predominant species were *B. ubonensis* (*n* = 252, 27.00% of isolates) and *B. multivorans* (*n* = 250, 26.8% of isolates), followed by *B. cenocepacia* (*n* = 175, 18.70% of isolates). Dataset 2 isolates were distributed across 29 different countries, with most isolates from Australia (*n* = 294) and the USA (*n* = 268) at 31.48% and 28.69% of genomes, respectively. The predominant isolate sources were North America (*n* = 426, 45.61%) followed by Oceania (*n* = 295, 31.58%) (Supplementary Table 3). In Dataset 2, Europe comprised a lower proportion of isolates than Dataset 1 at 5.57% (*n* = 52). In both Dataset 1 and Dataset 2, the main source of isolation was from people with CF at 46.5% and 45.70%, respectively. There was a similar number of environmental isolates observed in both datasets (32.4% and 39.9% of isolates). This corroborates their ubiquitous nature in the environment, but also the ability to infect persons with CF.

The average genome lengths of Dataset 1 and Dataset 2 were 7.27 Mb and 7.36 Mb, respectively. The number of contigs in Dataset 1 ranged from 1 to 7 contigs (average = 3.53 contigs) compared to a contig range of 1 to 198 (average = 80.48 contigs) in Dataset 2 (Supplementary Figure 2). The lower contig range in Dataset 1 reflected that of only complete Bcc genomes, whereas Dataset 2 included both complete and draft genomes. The average N50 values and GC content (%) for the assembled genomes were similar for both datasets. The N50 values were 3.29 x10^6^ and 3.72 x10^6^, whilst the average GC (%) content was 66.88% and 67.01% for Dataset 1 and Dataset 2, respectively (Supplementary Figure 2). The genomic features of the datasets mimic that of previously published Bcc genomes and their large, multi-replicon structures.

### Updating of Clonal Complexes in the PubMLST database

As there can be discrepancies between MLST profiles based on the sequencing and MLST data derived from Sanger sequencing and human errors during PCR amplification or data upload (Peeters *et al*., 2020), the ccs within the Bcc database were reviewed as part of this work. For instance, the coverage of Illumina sequencing can uncover genomic variants that are missed by Sanger sequencing (Peeters *et al*., 2020). Originally, there were 64 ccs defined within the PubMLST database at the time of analysis (on 26^th^ January 2026). These 64 ccs were assessed for the number of associated STs, based on the criteria of ≥5 loci matches. There were 23/64 ccs (35.94%) that had at least 10 isolates in the database.

All STs and respective loci (*n* = 2,517) were extracted from the PubMLST Bcc typing database and used for an input to Phyloviz (Nascimento *et al*., 2016). Initial clustering of groups was performed using a full MST goeBURST, which identified 106 clusters with a central ST (Supplementary Table 1; Supplementary Figure 1). These 106 clusters were cross-checked for suitability using BURST analysis on PubMLST. If a central ST was not identified using both methods, the cc was not defined in the database. Overall, there were 21 ccs designated on the PubMLST platform (Jolley *et al*., 2018) based on meeting the following criteria: (i) the observation of a central ST using both Phyloviz and PubMLST and (ii) having ≥10 isolates in the database associated with that central ST (Supplementary Table 1). When updating the ccs within PubMLST, conscious efforts were used to ensure that complexes well defined in the literature, such as the ST-31 in *B. cenocepacia* (Saroha et al., 2023) and the ST-15 in *B. multivorans* (Baldwin et al., 2007) retained the original cc nomenclature, irrespective of the central ST designation.

### The Bcc cgMLST scheme consists of 1,925 loci that span a variety of functions

PIRATE (Bayliss *et al*., 2019) and PEPPAN (Zhou *et al*., 2020) pangenome alignment tools were used to identify 30,590 and 41,796 total genes and 3,456 and 3,467 core genes, respectively, within Dataset 1 (*n =* 142 genomes) at the 95% core gene threshold. The total genes were consolidated between the two tools, identifying a consensus of 3,385 core genes. Quality filtering was performed on the consensus core genes by identifying presence in the larger validation Dataset 2 (*n* = 934 genomes), and filtering for invalid alleles when comparing to the PubMLST database (Jolley *et al*., 2018). This removed genes from the dataset that were either (i) absent in PubMLST after curation or (ii) genes that were off the end of contigs, reducing the number of core genes to 2,170 at the 95% threshold. To ensure that the scheme was robust and genomes could be automatically annotated in the PubMLST database (Jolley *et al*., 2018), a stricter 99% core gene threshold was ultimately chosen using the genomes in Dataset 2. The final Bcc cgMLST scheme consists of 1,925 loci, which have been assigned an arbitrary ‘BCC’ number for identification. At the time of writing (29^th^ April 2026), 1,465 of 1,889 (77.6%) Bcc genomes have been assigned a cgST on PubMLST (Jolley *et al*., 2018).

The core genes within the cgMLST scheme were assessed for the number of allelic variants and respective loci lengths using a previously published Python script (https://github.com/artmisk13/cgmlst_hinf/blob/code/core_gene_allelic_variability.py) (Krisna *et al*., 2024). The number of alleles ranged from 75 (Bcc12695) to 899 (Bcc01520), with a median and mean average of 539 and 508 allelic variants, respectively. Allele lengths ranged from 138bp (Bcc19985) to 4,842bp (Bcc1520), with a median allele length of 855bp (Supplementary Figure 3). A total of 1,839 (95.5%) of the core genes in the cgMLST scheme were provided with a putative COG annotation using eggNOG mapper (Cantalapiedra *et al*., 2021). There were 112 (6.1%) instances where more than one COG category was assigned to a locus. Where multiple COG categories were assigned, these were split into singular categories and the total numbers tallied for each category. In all, the core genes were assigned to 20 COG categories, comprising 4 KEGG (Kanehisa and Goto, 2000; Kanehisa *et al*., 2016) general definition groups. Most annotations were assigned to the ‘S’ category of unknown function (*n* = 439 core genes, 23.9%). Only one locus (Bcc0465, *hdaH*) was annotated with ‘B’ for Chromatin structure and dynamics. This locus was also assigned ‘Q’ for secondary metabolites biosynthesis, transport, and catabolism (Supplementary Figure 4; Supplementary Table 4).

### The cgMLST core gene clustering provides a more in-depth, hierarchical understanding of Bcc epidemiology compared to established methods

There were 908 unique cgSTs assigned to 913 (97.75%) of genomes in the validation dataset (Supplementary Table 3). To assess similarity of the isolates, core gene clusters (cgc) were used to group the cgSTs using different thresholds. This is a single-linkage clustering method that matches isolates based on a defined maximum number of allelic differences. In the Bcc cgMLST scheme, cgc groups of 1,750, 1,000, 500, 200, 100, 50, and 25 allelic mismatches were used. The lower allelic mismatch thresholds provide a high-resolution analysis of the Bcc isolates.

When mapping metadata onto a phylogeny built with cgMLST, the resolution of clusters was more evident compared to MLST as expected. However, no immediate visual correlations could be inferred from the linking of metadata to cgcs. At the cgc1750 level, concordance occurs with most Bcc species. An exception is *B. ubonensis* that exhibits at least 10 distinct clusters. Within *B. multivorans*, most (56%; *n =* 139) isolates are grouped in one cluster at cgc1750, with multiple alternative clustering groups (Figure 2A).

**Figure 2.**
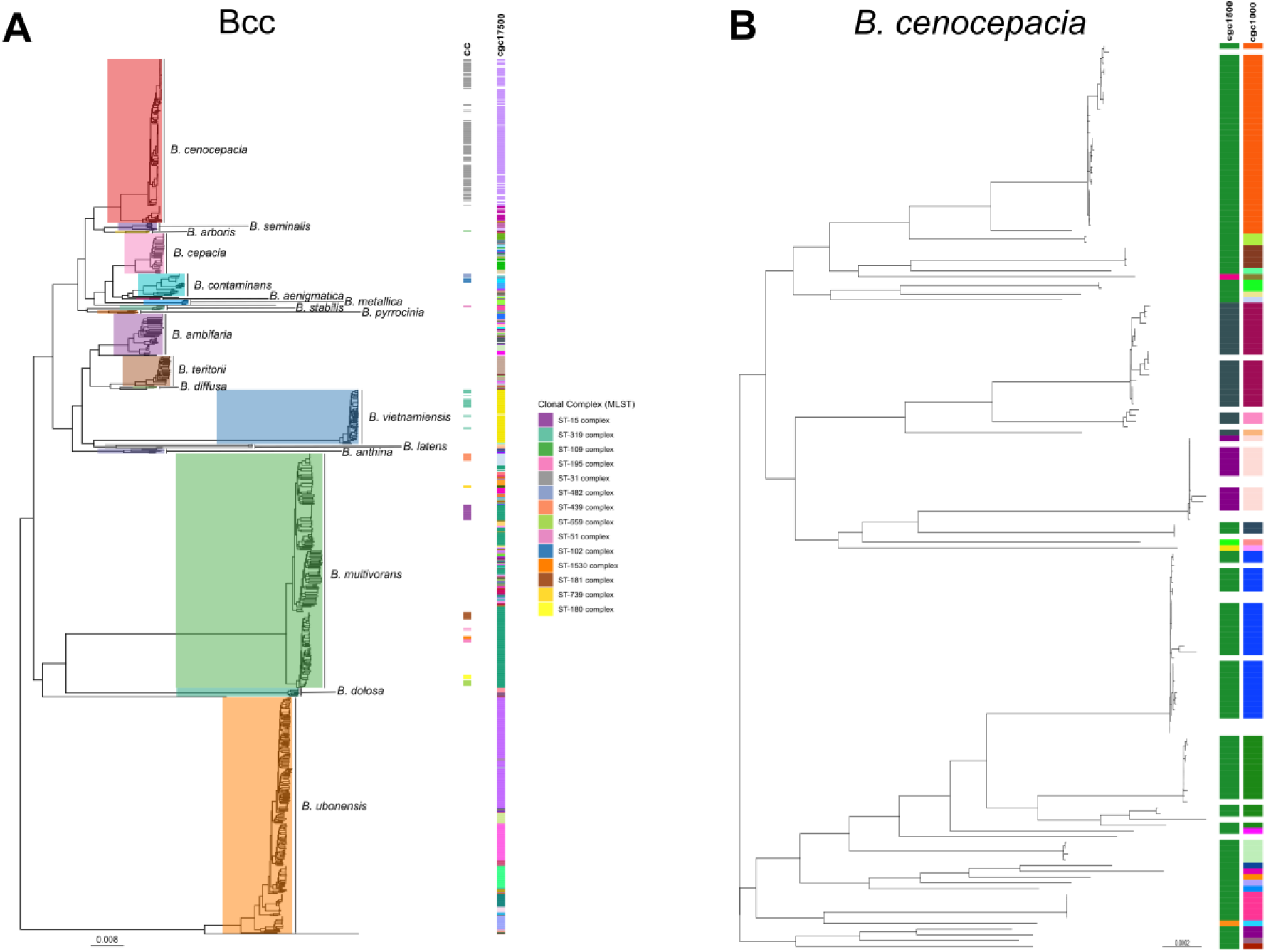
| Assessment of the BCC population structure to identify discontinuities using ReporTree. Stable regions (blocks) are shown compared to the number of cgMLST allelic mismatches. The top 8 allelic mismatch thresholds, identified using ReporTree, are shown using dashed lines. These represent 1891, 1834, 1805, 1639, 1407, 831, 120, and 58 loci mismatches.

**Figure 3.**
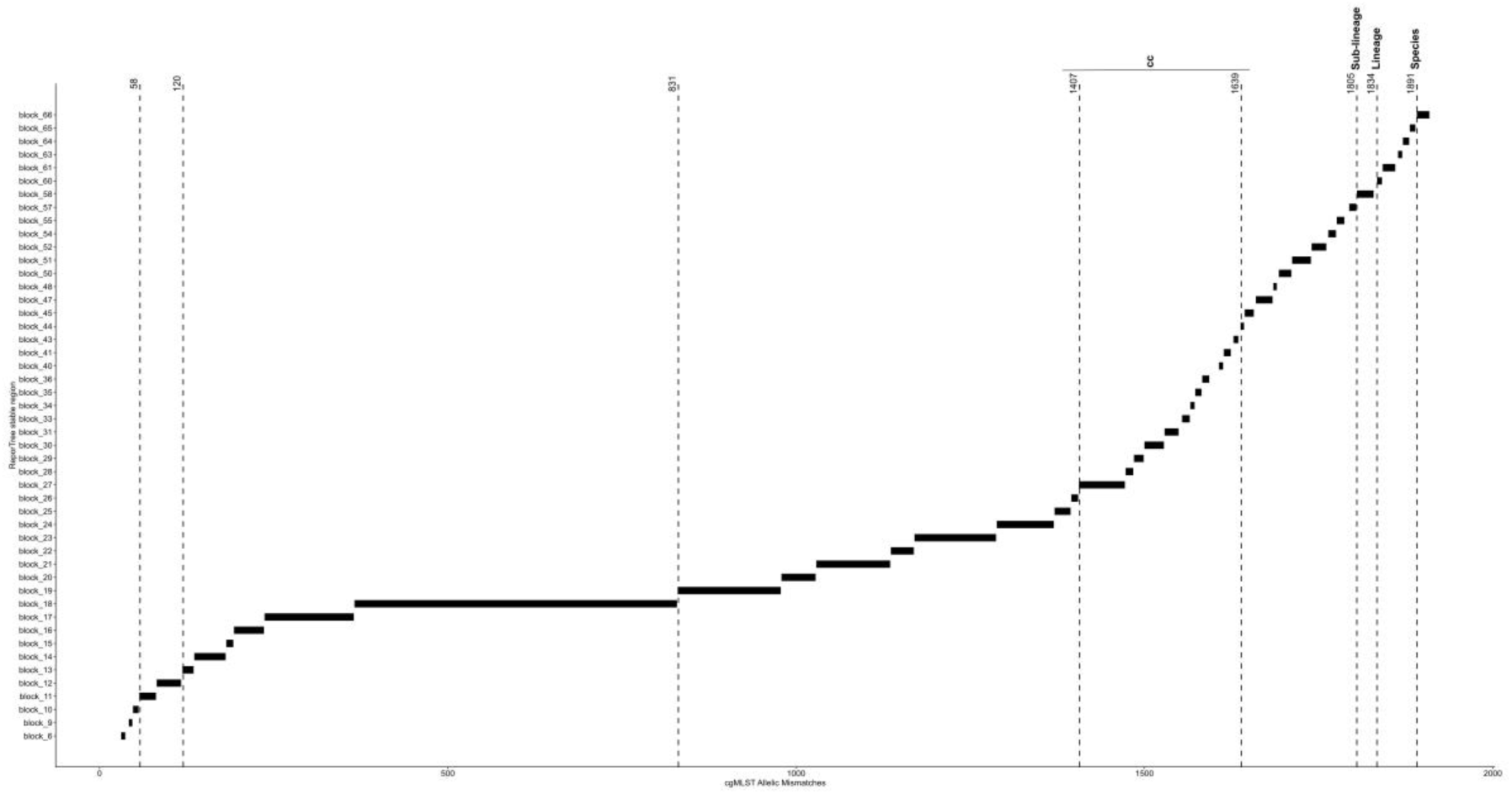
| (A) The population structure of 934 BCC genomes from the validation dataset. Maximum Liklihood phylogenetic trees have been built using IQ-Tree with 1,000 bootstraps, using the core gene loci from PubMLST. Metadata shows the clonal complexes and top-level cgc17500 clustering (B) A subset of the ML tree, detailing the B. cenocepacia (n = 175) population in high resolution. Metadata shows the clustering at cgc1500 and cgc1000, providing a high-resolution analysis of the B. cenocepacia population structure.

To illustrate the importance of cgc for epidemiological purposes, *B. cenocepacia* (*n* = 175) was used as an example. The majority of *B. cenocepacia* isolates (*n* = 107; 61.14%) belonged to a single cc (ST-31 complex). Using cgMLST clustering, *B. cenocepacia* could be divided into at least 7 and 29 subgroups at cgc1500 and cgc1000, respectively (Figure 2B). The ST-31 complex specifically was clustered into 4 and 15 groups at the cgc1500 and cgc1000 thresholds. One of the 7 *B. cenocepacia* cgc1500 clusters was observed in two phylogenetic lineages. This cluster was diverged into distinct clusters at the cgc1000 threshold, correlating with the phylogenetic groups (Figure 2B).

### Defining allelic mismatch thresholds within the Bcc population

The Bcc population structure was investigated for discontinuities using ReporTree (Mixão *et al*., 2023); a tool that identifies hierarchal genetic clusters which can be used for epidemiological tool development and analysis. There were 155 unique clusters, which are made up of isolate groups at one specific allelic mismatch threshold, that were defined in the Bcc population. Of this, there were 48 stable regions (blocks) where the clustering composition between groups was uniform or had little variation (Supplementary Table 5; Figure 3). These clustering regions were compared to the respective Adjusted Rand Index and Adjusted Wallace scores to assess optimality (Supplementary Table 5). A total of 14 allelic mismatch thresholds were chosen to represent the overall population structure. The first 8 thresholds were defined based on the clustering output from ReporTree (Mixão *et al*., 2023) (Figure 3). The first three thresholds were between 34-120 locus differences from the maximum 1,925 loci in the cgMLST schema, corresponding to the fact that most discontinuity within the population occurred in these higher boundaries. An allelic mismatch threshold at 1891 locus differences (98.23%) was chosen to represent a ‘species-level’ boundary (Figure 4). There were 43 unique LIN codes that represented 23 species at this boundary. At this level, LIN codes corroborated the distinction of *B. orbicola,* formally known as *B. cenocepacia* IIIB (designated LIN prefix 16), from *B. cenocepacia* IIIA (LIN prefix 17) (Morales-Ruíz et al., 2022). PubMLST isolates belonging to the *B. cenocepacia* IIIC and IIID lineages (Vandamme *et al*., 2003) were also designated prefixes of ‘15’ and ‘27’, respectively, highlighting the ability of LIN codes to group isolates at the level of *B. cenocepacia recA* linages. A second threshold was then defined at 1,834 locus mismatches (given the boundary of “lineage” level). ‘Sublineage’ was defined using a boundary of 1,805 locus mismatches, which is used to further distinguish between lineages that share a common ancestor.

**Figure 4.**
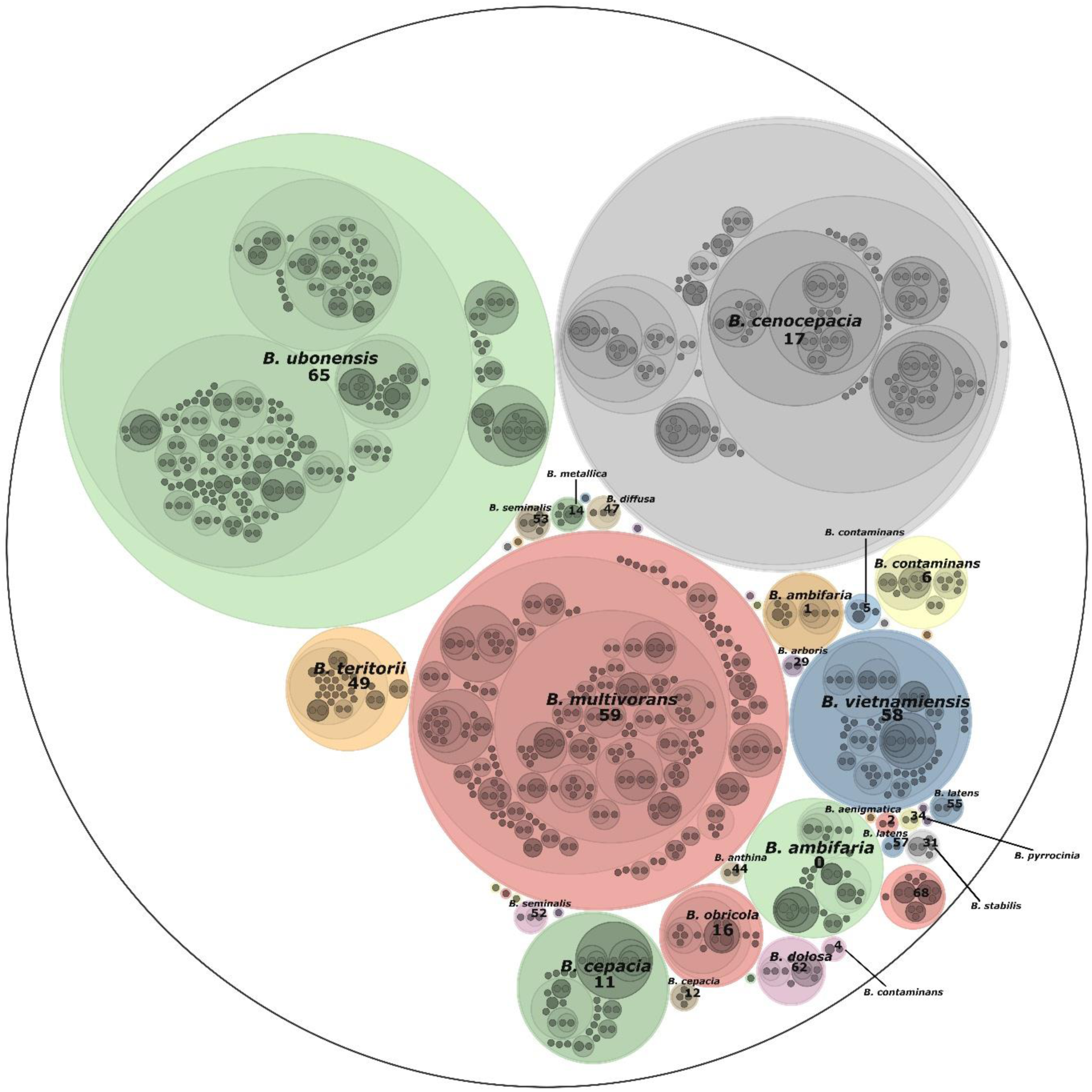
| The hierarchical structure of the BCC population (n = 934) based on LIN codes, with the top threshold (1,891 cgMLST loci mismatches) annotated. Each isolate has a designated 14-threshold LIN code, which is nested within the parental prefix. This is illustrated by the nested circles within each species. The size of the segments illustrates the number of validation genomes used for that species. Figure was created using LINviz on PubMLST.

The 4^th^ and 5^th^ LIN thresholds (1639 and 1407 locus mismatches, respectively) corresponded to the clonal complexes within the Bcc (Table 2). *B. multivorans* had the most diverse range of associated LIN codes, of which 5/8 (62.5%) ccs diverged at the 5^th^ threshold. The ST-195 complex, within the *B. multivorans* population, had two associated LIN codes (59_0_0_0_17 and 59_0_0_0_18), highlighting a sub-structure within this cc. Whilst inferences between cc and LIN code can be made, there are limitations to this due to the small numbers of cc annotations in the PubMLST Bcc database. For instance, of the 934 genomes in Dataset 2, only 192 (20.6%) had an annotated cc.

**Table 2.** | Comparison of the LIN prefixes to clonal complex (CC) and *Burkholderia* spp.

| **Species** | **Clonal Complex** | **LIN Prefix** |
|---|---|---|
| *Burkholderia arboris* | ST-109 complex | 29_0_0_0 |
| *Burkholderia cenocepacia* | ST-31 complex | 17_0_0_0 |
| *Burkholderia contaminans* | ST-102 complex | 6_1_0_2 |
| *Burkholderia contaminans* | ST-482 complex | 6_1_0_1 |
| *Burkholderia multivorans* | ST-15 complex | 59_0_0_11 |
| *Burkholderia multivorans* | ST-1530 complex | 59_0_0_0_21 |
| *Burkholderia multivorans* | ST-180 complex | 59_0_0_0_27 |
| *Burkholderia multivorans* | ST-181 complex | 59_0_0_0_16 |
| *Burkholderia multivorans* | ST-195 complex | 59_0_0_0_17 & 59_0_0_0_18 |
| *Burkholderia multivorans* | ST-439 complex | 59_0_0_33 |
| *Burkholderia multivorans* | ST-659 complex | 59_0_0_0_14 |
| *Burkholderia multivorans* | ST-739 complex | 59_0_0_26 |
| *Burkholderia stabilis* | ST-51 complex | 31_0_0_0 |
| *Burkholderia vietnamiensis* | ST-319 complex | 58_0_0_0 |

The lowest 6 mismatch thresholds (14, 7, 4, 2, 1, and 0 locus differences) were chosen to facilitate epidemiological studies, with high-resolution taxonomic clusters at these boundaries. The lowest threshold (0 allelic mismatches) is used for indistinguishable isolates that only differ by missing cgMLST loci. These thresholds were chosen through an arbitrary means, rather than hierarchical clustering as before, and will be used to identify transmission events for use in outbreak analysis.

The 14-LIN thresholds were added to the Bcc PubMLST database (Jolley *et al*., 2018), with a taxonomy defined for each genome that harboured a cgST. Overall, there were 67 unique LIN codes assigned to 1,309 distinct Bcc cgSTs on PubMLST (17^th^ March 2026). This encompassed the annotation of 86.9% (1,217/1,401) of genomes in the Bcc database. There were 71 unique LIN codes associated with species, 102 with lineages, and 162 with sublineages. At the 1-locus mismatch clonal group level (13^th^ threshold), used for molecular epidemiology, there were 1,253 unique LIN codes.

### Application of LIN codes to published Bcc datasets

The Bcc LIN codes were applied to three published Bcc datasets to illustrate their use in genomic epidemiology and outbreak surveillance. In the first example, LIN codes were used to identify genetic differences between two distinct Bcc species, *B. arboris* and *B. cepacia*, part of a healthcare-associated outbreak in Germany (Bender *et al*., 2022). It was previously observed, using an *ad hoc* cgMLST scheme, that these two species were associated with mouthwash contamination in multiple hospitals (Bender *et al*., 2022). Our LIN codes were able to further differentiate differences within the two species clusters. There were 9 of 11 (81.8%) *B. cepacia* isolates that harboured identical LIN codes, with the 2 other isolates differing at the 13^th^ threshold (1 locus difference) (Figure 5). In addition, the unrelated isolate (761-18) from hospital B was designated as *B. contaminans* using LIN codes, rather than the designation of *B. cepacia* as per the outbreak publication.

**Figure 5.**
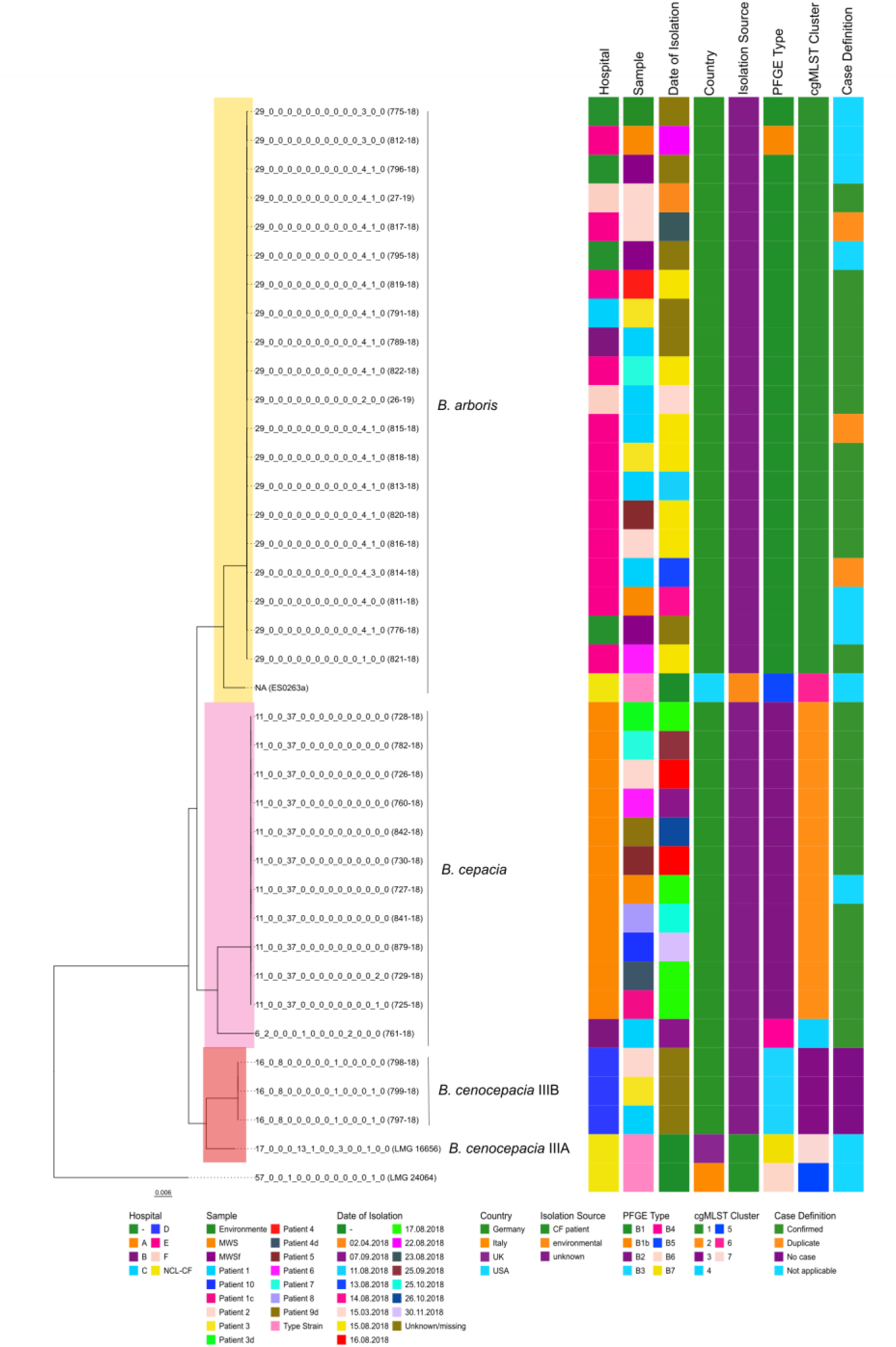
| Maximum Likelihood phylogeny of 38 BCC genomes (n = 12 of B. cepacia, n = 21 of B. arboris, n = 4 of B. cenocepacia, and n = 1 of B. latens), built using RAxML with 100 bootstraps with the cgMLST loci. Recombination has been adjusted for using ClonalFrameML. This is a re-analysis of a published outbreak of B. cepacia and B. arboris in contaminated mouthwash by Bender et al., (2022). One isolate (761-18) has been designated as B. contaminans using the LIN codes, opposed to the original designation of B. cepacia in the original publication. The phylogenetic tree has been rooted on B. latens LMG 24064 (PubMLST ID 404). Associated outbreak metadata has been used to supplement the full 14-threshold LIN codes.

Within the *B. arboris* cluster (Bender *et al*., 2022), 13 of 21 (61.9%) isolates harboured indistinguishable LIN codes. The remaining 4 isolates differed at the 12^th^ threshold (2 locus differences). Despite differing at this threshold from other isolates (Bender *et al*., 2022), isolates 812-18 (PubMLST ID: 5591; mouthwash solution isolate) and 775-18 (PubMLST ID: 5581; filling patient isolate) were identical (Figure 5). This confirmed that the two batches of mouthwash solution were likely contaminated by the same *B. arboris* isolate.

Analysis of a *B. contaminans* outbreak in an eye clinic in Norway (Spilker *et al*., 2022) showed that LIN codes could cluster epidemiologically related isolates at the fine thresholds. The LIN codes showed that 12 *B. contaminans* isolates from Norway, that were associated with either the clinic ventilation duct or endophthalmitis patients, had indistinguishable barcodes. Two isolates from the clinic ventilation duct (HI4951 and HI4957) were distinguished from this cluster at the 12^th^ (2 locus differences) and 13th thresholds (1 locus difference), respectively (Figure 6). This highlights the concordance of LIN codes with genomic analysis using single nucleotide polymorphisms (SNPs).

**Figure 6.**
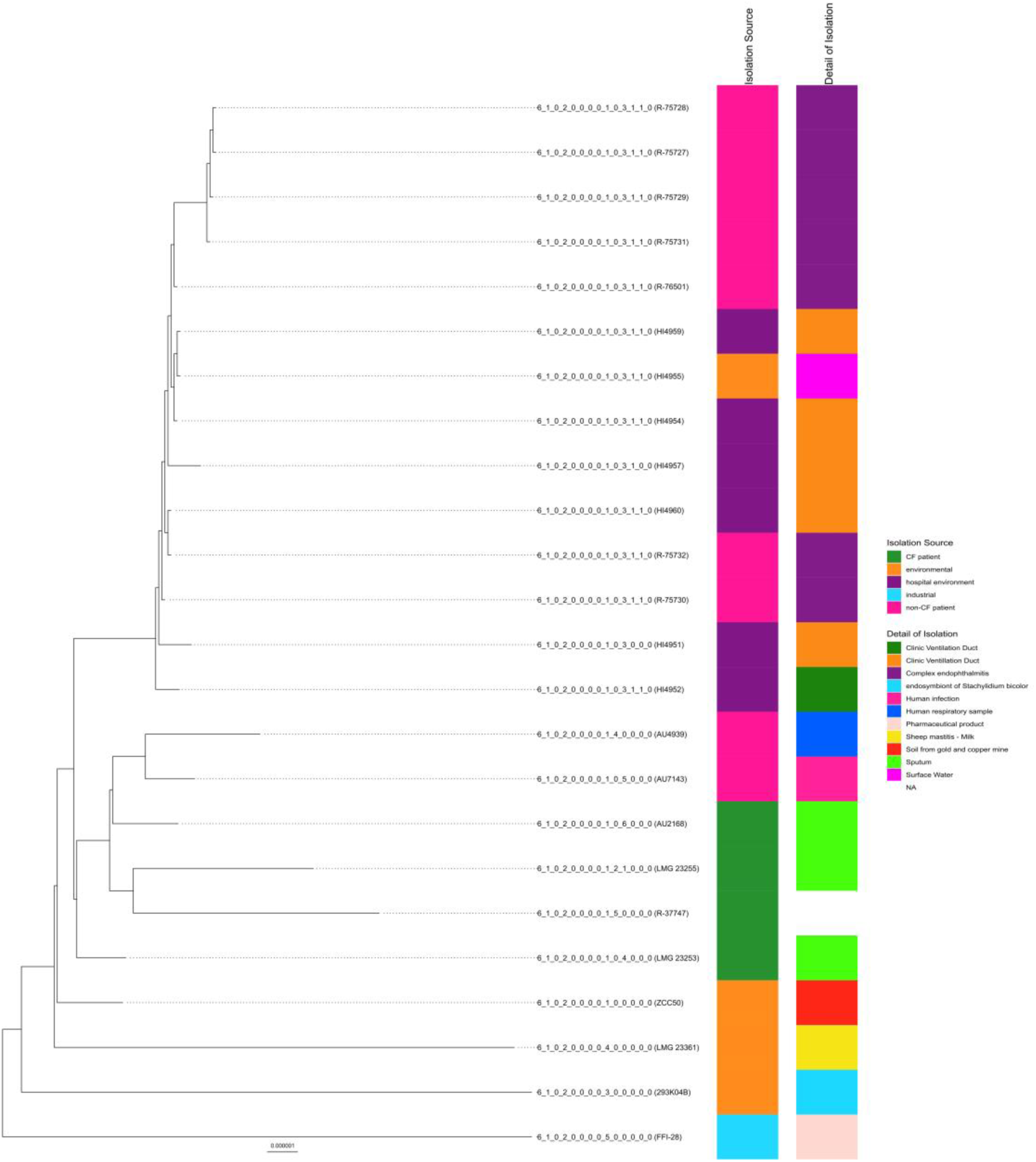
| Maximum Likelihood phylogeny of 24 B. contaminans genomes, built using RAxML with 100 bootstraps with the cgMLST loci. Recombination has been adjusted for using ClonalFrameML. This is a re-analysis of a published outbreak of endophthalmitis which was traced to a clinic ventilation system following cataract surgery Spilker et al., (2021). The phylogenetic tree has been midpoint rooted for analysis. Associated outbreak metadata has been used to supplement the full 14-threshold LIN codes.

LIN codes were also used to reproduce analysis of a *B. multivorans* isolate panel (*n* = 77 isolates) (Parfitt *et al*., 2023). This increased the resolution regarding the genetic diversity of *B. multivorans* isolates within the previously defined lineages. The LIN codes recaptured the closer relatedness of lineage 1 (*n* = 27) compared to lineage 2 (*n* = 49) but further identified subgroups within the lineages. Five isolates in lineage 2 were not designated a cgST or LIN code due to not meeting the threshold criteria, so were excluded from enumeration. The majority of lineage 1 isolates (*n* = 23; 85.1%) differentiated at the 5^th^ threshold (1,407 locus differences), whereas *n* = 17 (38.6%) and *n =* 27 (61.4%) of lineage 2 isolates differed at the 3^rd^ and 4^th^ threshold, respectively (Figure 7). Differentiation within lineages could be important for the elucidation of both global epidemiology of Bcc species, but also for identifying and monitoring the emergence of variants of public health concern.

**Figure 7.**
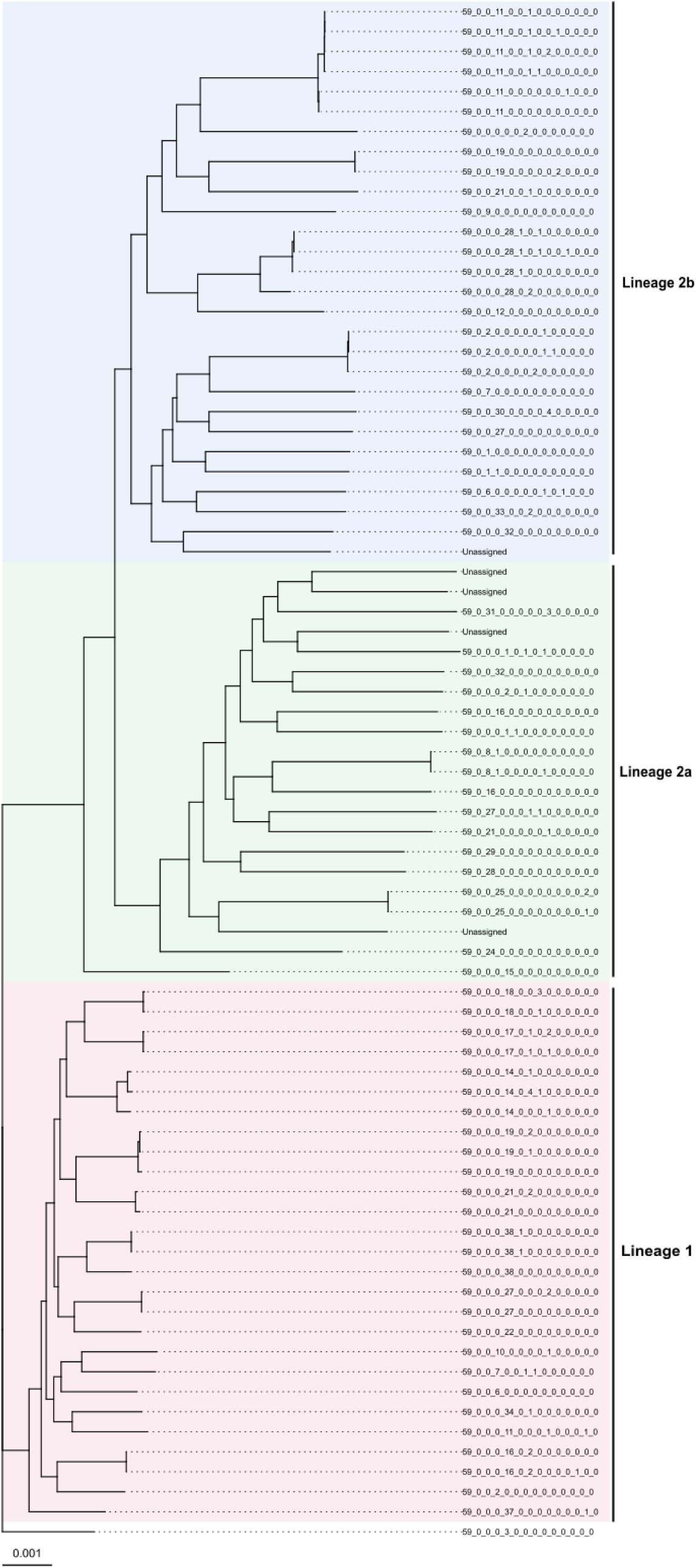
| Maximum Likelihood phylogeny of 77 B. multivorans genomes, built using RAxML with 100 bootstraps with the cgMLST loci. Recombination has been adjusted for using ClonalFrameML. This is a re-analysis of the B. multivorans population previously published by Parfitt et al., (2023). The phylogenetic lineages (1, 2a, and 2b) are defined and compared to the full 14-threshold LIN code.

## Discussion

The genomic population biology of the Bcc is still subject to much exploration, with only a limited number of species studied in-depth (Parfitt *et al*., 2023). An increase in the understanding of Bcc diversity is key to aiding epidemiology, pathology and disease surveillance. Whilst global Bcc infection incidence is low, this group of bacteria are challenging to identify and diagnose, as well as difficult to treat by nature. This has a profound effect on morbidity and mortality of immunocompromised individuals, such as those with CF (Salsgiver *et al*., 2016; Kenna *et al*., 2017). Despite advances in infection control to reduce overall Bcc prevalence and transmission in CF (France *et al*., 2008; Salsgiver *et al*., 2016), a lack of knowledge for non-CF infection and its control can lead to outbreaks. Combined, this presents a global public health threat for the Bcc, so early detection and treatment are key. We have built on the current molecular methods, such as conventional MLST (Baldwin *et al*., 2005) and extended MLST schemes (Spilker *et al*., 2009), for Bcc detection by increasing the resolution required for fine discrimination in epidemiological studies. This work presents a standardised 1,925 loci cgMLST scheme and subsequent 14-threshold LIN code nomenclature to describe the Bcc population.

Even though the current analysis was developed on a subset of 19 Bcc species, the resulting cgMLST and LIN codes proved to be applicable to all currently described Bcc species (Figure 4). Application across Bcc species is important due to the redefining of taxonomic groupings within the complex, with new species still being defined, such as *B. orbicola* (Morales-Ruíz *et al*., 2022). It was important to develop a cgMLST and LIN code scheme for the Bcc, instead of one for each single species, due to isolate identification in clinical laboratories. Matrix-assisted laser desorption ionization-time of flight Mass Spectrometry (MALDI-ToF MS) can accurately identify Bcc isolates to the genus level, but species-level identification is not always accurate using this method (Fehlberg *et al*., 2013). Therefore, the use of WGS can immediately provide lineage-level identification through upload to the PubMLST platform, enabling assignment of a cgST and LIN code.

MLST has been useful for elucidating the Bcc population (Baldwin *et al*., 2005), including re-classification of isolates and identifying new species (Spilker *et al*., 2009). Due to their slow molecular clock, the conserved seven MLST housekeeping genes have also been shown to remain stable during chronic infection (Waine *et al*., 2007). However, the Bcc are highly recombinant which makes it difficult to capture finer differences between isolates using smaller numbers of loci. For instance, horizontal gene transfer (HGT) accounts for around 10% of the genetic material in Bcc genomes (Sousa *et al*., 2011). HGT contributes for the genetic variation between Bcc lineages that contribute to niche adaptation, such as within *B. cenocepacia* J2315, where the acquisition of 14 genomic islands (Holden *et al*., 2009) facilitates survival and pathogenesis in the CF lung (Sousa *et al*., 2011). LIN codes can help to capture the genetic variation between Bcc lineages, which in turn could identify clusters that have adapted to cause endemic lineages. For instance, if a particular LIN cluster is detected over time across epidemiologically linked cases, it can be assumed that lineage possesses genetic characteristics that favour persistence, transmission, or niche adaptation. This ultimately provides a framework for investigating evolutionary processes of Bcc lineages that become established and persist within a population.

Despite strict and effective infection control measures to reduce Bcc transmission between CF individuals, particularly in the case of epidemic CF lineages (Saiman and Siegel, 2004), acquisition from environmental sources remains a contributing factor to infection (Baldwin *et al*., 2007; Drevinek and Mahenthiralingam, 2010). Antibiotic treatment is used to combat Bcc infection but remains challenging due to the intrinsic and acquired multi-drug-resistant nature of this bacterial group. For instance, over 50% of individuals with CF have shown resistance to the10 most used antibiotic therapies (Aaron *et al*., 2000). Beyond this, the Bcc also exhibit resistance to biocides, and antiseptics through mechanisms such as efflux pumps (Rushton *et al*., 2013; Ahn *et al*., 2016). Understanding links between bacterial populations and antimicrobial resistance characteristics are important for outbreak detection and control (Maiden *et al*., 2013). LIN codes have previously been shown to identify associations between lineages and AMR in other pathogens (Unitt *et al*., 2025), as well as the linkage of virulence-associated allelic variants (Delgado-Blas *et al*., 2025), which could be applied in the future to the Bcc.

The Bcc LIN codes also proved effective for epidemiological tracking at the species and strain level, as illustrated by the meta-analysis of three published datasets herein (Table 1). Fine levels of resolution can be observed in the Bcc when using genetic clusters between 14 and 0 allelic mismatches. For example, the use of LIN codes highlighted genetic differences and 1 and 2 locus mismatches for the *B. cepacia* and *B. arboris* clusters, respectively, associated with an outbreak in Germany linked to mouthwash solution (Bender *et al*., 2022). We also highlight the congruence of the LIN codes to SNP analysis when assessing a *B. contaminans* outbreak (Spilker *et al*., 2022). Whilst SNPs may show further differences between the isolates due to analysing the whole genome instead of just the core, the use of LIN codes is the optimal choice for rapid detection of clusters or emerging lineages in an outbreak situation. Furthermore, only a genome is required to be uploaded to the PubMLST database for LIN code assignment, which is based on the genomes that are already present. This contrasts with SNP analysis which requires re-analysis to be performed with the inclusion of new genomes. This would in turn facilitate an ‘alert’ for public health officials so that they can actively respond in a timely manner. The incorporation of LIN codes into PubMLST creates this framework for real-time international surveillance, including the backwards compatibility with the MLST scheme. As new genomes are deposited, they can be assigned immediately to existing Bcc species, lineages and outbreak-associated groups without need for research or redefining historical nomenclature.

## Conclusions

Overall, we have established the first, unified, Bcc-wide cgMLST and LIN-code framework that delivers stable, taxonomy-inclusive identifiers accurate to the isolate lineage level. The LIN codes provide a portable, publicly available advancement in the rapid detection of concerning or emerging Bcc lineages and outbreak scenarios. These provide a computer readable schema, which are backwards compatible to nomenclature such as 7-loci based STs and ccs. This LIN code scheme this will enhance global public health and Bcc disease surveillance through the monitoring of disease prevalence and endemic lineages.

## Author Contributions

The CReDIT contributor roles taxonomy was used to recognise author contributions. Conceptualization: KMP and MCJM. Methodology: KMP, MAK and KAJ. Software: KMP, MAK, and KAJ. Validation: KMP. Formal analysis: KMP. Investigation: KMP. Resources: KAJ and MCJM. Data curation: KMP, KAJ and CP. Writing – original draft preparation: KMP. Writing – review and editing: All authors. Visualisation: KMP, EM, and CP. Supervision: MCJM. Project administration: KMP and MCJM. Funding acquisition: MCJM.

## Notes

### Competing Interest Statement

The authors have declared no competing interest.

