## Supplementary figures and images for "Development of a core genome Multilocus Sequence Typing (cgMLST) and Life Identification Number (LIN) code schema for the *Burkholderia cepacia* complex"

### Supplementary Figure 1

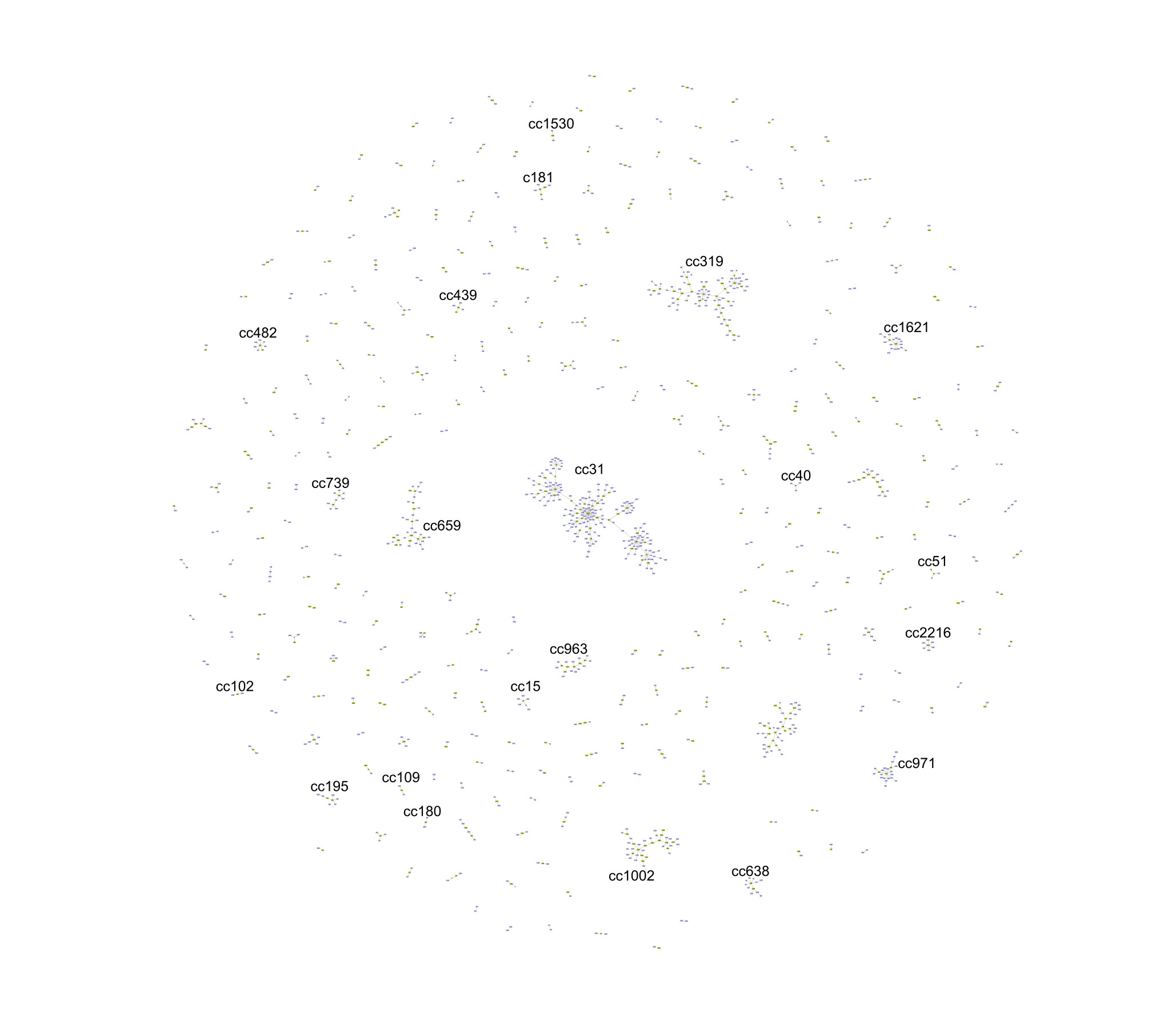

### Supplementary Figure 2

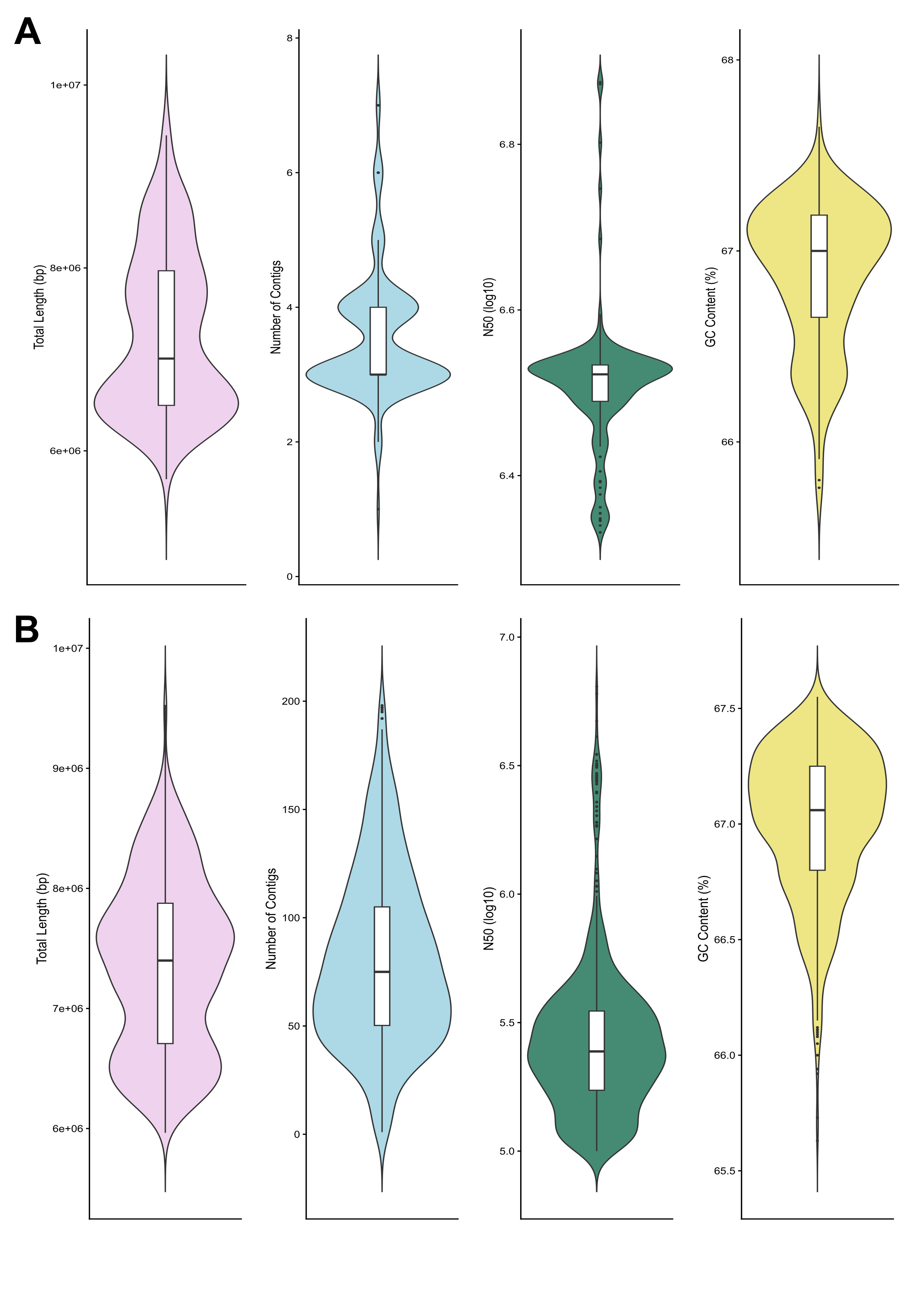

### Supplementary Figure 3

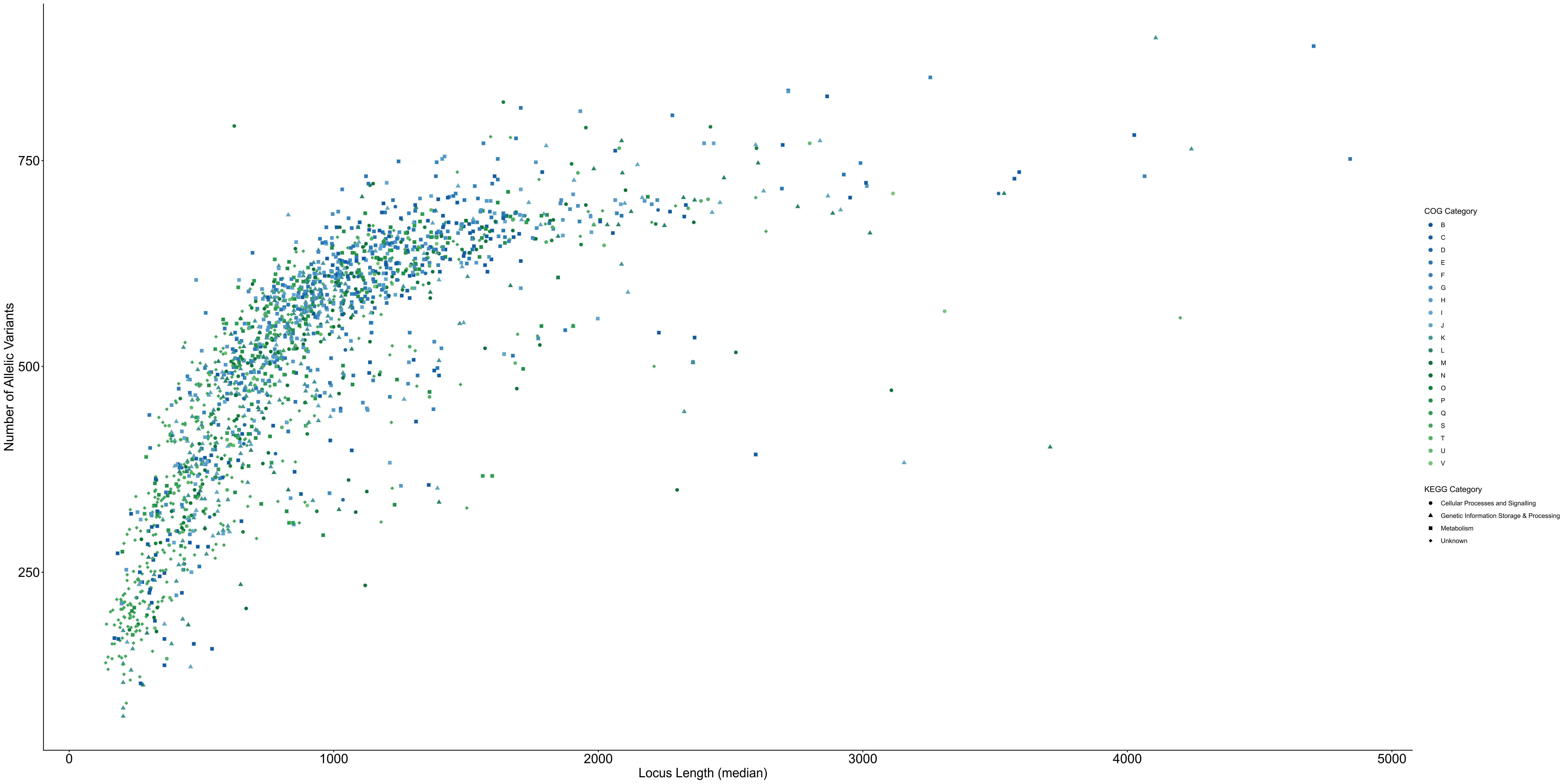

### Supplementary Figure 4

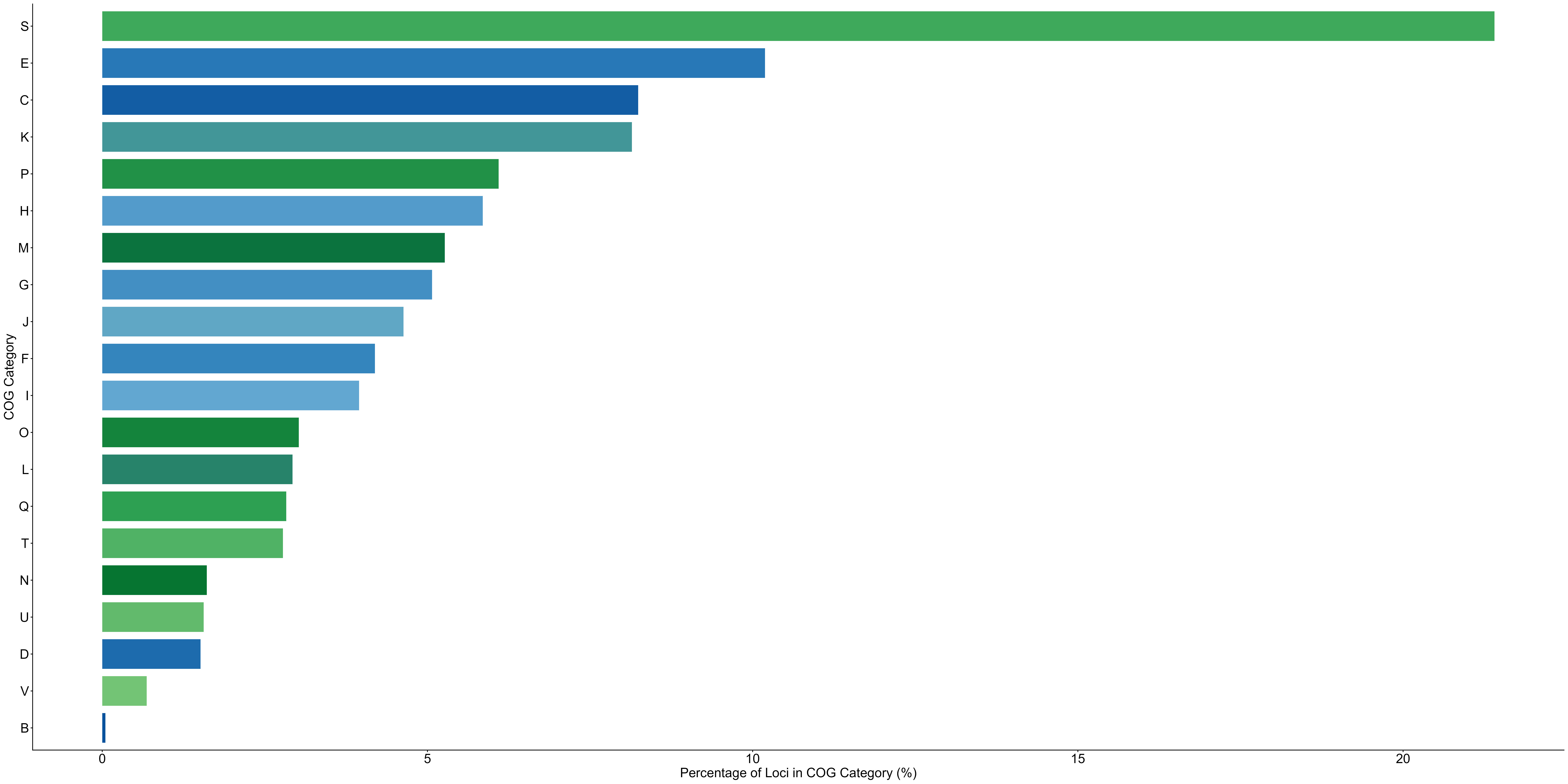
